# The Sweet Spot of Plasticity in Perceptual Learning Depends on Training Variability

**DOI:** 10.64898/2025.12.17.694881

**Authors:** Giorgio L. Manenti, Caspar M. Schwiedrzik

## Abstract

Perceptual learning improves sensory discrimination, yet the brain must balance specificity and generalization, especially in variable environments. To investigate how variability shapes perceptual learning, we used functional magnetic resonance imaging while human subjects performed an orientation discrimination task with low or high task-irrelevant variability in spatial frequency. Behaviorally, both training regimes improved performance, but only high variability training enabled transfer to a novel location. Retinotopic orientation representations in early visual areas changed similarly across groups, indicating that variability does not alter local sensory representations. In contrast, functional connectivity differed: low variability training enhanced task-dependent coupling between the intraparietal sulcus and V2, while high variability training differentially affected connectivity with hV4. These results support a “sweet spot” model of perceptual learning, in which the locus of learning flexibly depends on the training environment. But rather than modifying sensory representations, variability reconfigures network-level readout, enabling specific or generalizable improvements in perception.

## INTRODUCTION

Perception is inherently adaptive, continuously adjusting to the demands of an ever-changing sensory world. A critical aspect of this adaptability is the capacity to generalize learning beyond the specific conditions under which it was acquired. Visual perceptual learning (PL), the experience-driven improvement of visual abilities, exemplifies this adaptability by enabling discrimination of subtle, even initially imperceptible stimulus differences ^1,2^. Paradoxically, however, PL is often strikingly *specific* to the trained stimuli or tasks ^3,4^, in contrast to many other forms of learning that generalize more broadly ^5^. This specificity suggests potential fundamental constraints on sensory plasticity and raises key questions about the mechanisms that govern how we can improve and/or recover our perceptual abilities.

Traditional theories attribute PL specificity to changes in early sensory representations, where neurons have small receptive fields and sharp feature tuning ^6–8^. Alternative accounts suggest that specificity may instead arise from how downstream circuits read out these highly specific early visual representations ^9,10^. While both frameworks can explain PL specificity, behavioral evidence shows that PL can generalize under certain conditions ^11–13^, including in patients with visual deficits undergoing PL-based rehabilitation ^14^. This challenges the view that PL originates solely in early visual areas. Understanding how the visual system allocates plasticity to support either specificity or generalization hence remains a central challenge for theories of PL.

One important variable that emerges as a potential regulator for determining specialized versus generalized PL is *variability* during training ^15^. Variability during training may encourage learners to extract rules or abstract principles that support performance beyond the specific training conditions. In contrast, training with uniform stimuli can drive PL toward specificity, reflecting an overfitting to the exact features of the training set. Notably, real-world visual environments are characterized by substantial variability ^16^, suggesting that mechanisms supporting generalization may have evolved under such conditions, whereas traditional laboratory training paradigms with very small stimulus sets represent an artificial situation that may favor specificity.

We have recently shown that systematically varying a task-irrelevant feature during training can be both highly effective in driving generalization as much as highly informative about the underlying neural mechanisms. For example, we could show behaviorally and in silico that task-irrelevant variability in spatial frequency (SF) or phase during orientation discrimination training enhances generalization to untrained SFs and locations ^12,17^. This is particularly striking because SF and location specificity are well-established phenomena in PL ^18–20^ and are often interpreted as direct reflections of receptive field properties of early visual areas thought to be involved in PL.

Conceptually, there are (at least) two computational strategies by which the visual system could support PL under stimulus variability ^12^: a specialization strategy recruits neurons narrowly tuned to the task-relevant feature *and* to co-varying stimulus properties - consistent with the classical view that plasticity occurs at the lowest cortical stage that differentially encodes the trained feature ^8^. Such highly selective representations offer high-fidelity discrimination, but their specificity can limit transfer. Conversely, a generalization strategy relies on broadly tuned, invariant representations in higher visual areas ^21^ that pool across input variability by means of their invariance properties. By responding consistently across variations in task-irrelevant features, these invariant neurons absorb trial-to-trial variability into their broad tuning, ensuring that the task-relevant information remains accessible even when the precise stimulus instantiation changes ^12^. This promotes transfer to novel stimuli yet may sacrifice precision relative to specialized early codes. These contrasting costs and benefits suggest that PL need not rely exclusively on one strategy; instead, where plasticity is expressed may depend on how the brain balances the variability and precision demands of the task at hand.

Building on these perspectives, we hypothesized that the brain flexibly selects between recruiting earlier (specialization) versus later (generalization) sensory neurons depending on the level of task-irrelevant variability during training. Therefore, rather than plasticity being fixed to an early sensory stage, as traditionally proposed ^8^, PL may emerge from a distinct physiological “sweet spot” across the sensory hierarchy that optimally matches the currently required tradeoff between precision and variability: training with high variability would favor recruitment of broadly tuned, invariant neurons that support generalization, whereas low variability training would engage more narrowly tuned populations to optimize precision ^12^.

To test how training dimensionality shapes perceptual learning, we trained human participants on an orientation discrimination task with either low or high task-irrelevant variability in SF while undergoing functional magnetic resonance imaging (fMRI). Orientation information is encoded redundantly across the ventral visual stream ^22^, and representations become increasingly invariant to SF at higher stages of the hierarchy ^23^. This provides a principled framework for identifying where plasticity emerges: if learning relies exclusively on specialized early mechanisms, training effects should localize to V1 or V2 ^24,25^; if, instead, task-irrelevant SF variability recruits SF-invariant representations, plasticity should shift to downstream ventral-stream regions like hV4. To adjudicate between these possibilities, we quantified representational changes at the retinotopic stimulus locations across the visual processing hierarchy using multivoxel pattern analysis (MVPA) and examined changes in functional connectivity between visual and parietal cortex as a proxy for readout. Parietal cortex is a central node in perceptual learning, implicated in linking sensory representations to decision making ^26,27^, and thus provides an entry point for assessing how learning-related signals propagate to downstream decision circuits.

Behaviorally, classical low-variability training produced the expected specificity, with learning failing to generalize to a new location. In contrast, high-variability training enabled robust generalization to an untrained location, replicating our previous findings ^12^. At the neural level, decoding accuracy across early visual areas increased with training, but in both groups, indicating that plasticity of early sensory representations alone cannot account for the divergent generalization profiles. Instead, learning regime-related differences emerged in connectivity patterns: training with low variability was associated with strengthened connectivity between V2 and IPS, whereas high task-irrelevant variability additionally modulated connectivity between hV4 and IPS. Notably, the hV4–IPS coupling reappeared during the transfer session, suggesting that interactions between higher-order visual cortex and parietal decision circuits play a key role in supporting generalization following training with task-irrelevant variability.

Together, these results motivate the view that the locus of plasticity in visual PL is not fixed, but instead depends on the structure of the training environment. Recognizing this flexibility reframes PL not as a property of a fixed cortical site, but as an emergent consequence of how the visual system integrates task demands with its representational hierarchy.

## RESULTS

### Behavioral learning and transfer effects

To investigate how variability during training affects generalization and where training-induced neural plasticity takes places, we trained 40 participants (23 female, 3 left-handed, mean age 27 yrs, SD 6.0 yrs) for four days on an orientation discrimination task (Fig. 1A) where they had to judge the tilt of a grating (clockwise or counterclockwise), placed 12.4° in the periphery in the lower right quadrant (see Methods). To-be-discriminated orientation differences ranged between 0.5° and 2.75°. Subjects were divided into two groups: the “low variability” group saw gratings with only one SF (1.70 cpd), while the “high variability” group saw gratings with three randomly intermixed SFs (0.53, 1.70, and 2.76 cpd). The SFs were chosen to minimize overlap between adjacent SF channels (see Methods).

**Figure 1:**
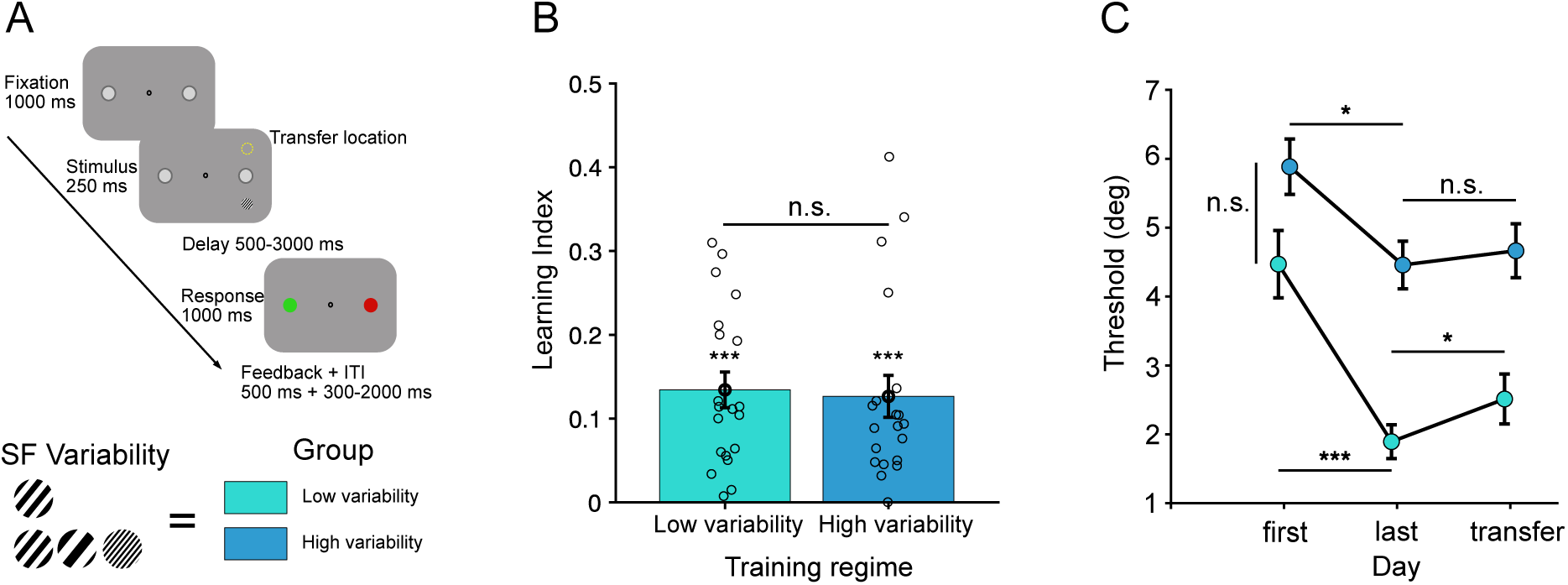
Experimental design, learning and generalization. **A**: Two groups were trained on an orientation discrimination task with either low SF (1.70 cpd) or high SF variability (0.53, 1.70, and 2.76 cpd). During training, stimuli were presented in the lower right quadrant, while the transfer location was in the upper right quadrant, 8° away. After a variable delay, participants indicated the side of a color cue corresponding to the stimulus orientation (green for clockwise). **B:** Learning in orientation discrimination, quantified as learning index (LI), was significant in all groups (low variability mean LI=0.13, permutation test, p<0.001, Hedges’ g=1.95; high variability mean LI = 0.13, permutation test, p<0.001, Hedges’ g=1.55). Task-irrelevant variability did not negatively affect the amount of learning (groups mean difference in LI=0.008, permutation test, p=0.815, Hedges’ g=0.07). Error bars reflect the standard error of the mean. **C**: We determined discrimination thresholds (using Weibull fits) per group and session. In the high-precision regime, both groups show significant learning effects on thresholds (low variability: mean [last-first]=−2.758, permutation test, p<0.001, Hedges’ g=−1.08; high variability: mean [last- first]=−1.43, permutation test, p=0.033, Hedges’ g=−0.67), which do not differ between groups (mean difference between groups [last-first]=1.15, permutation test, p=0.197, Hedges’ g=0.43). Furthermore, the low-variability training regime shows specificity of learning, as its threshold increases significantly for the location transfer (mean [transfer-last]=0.62, permutation test, p=0.039, Hedges’ g=1.08). For high variability, we find no significant change in thresholds at the new location (mean [transfer-last]=0.21, permutation test, p=0.361, Hedges’ g=0.09), suggesting generalization. Error bars reflect the standard error of the mean, corrected for between-subject variability ^29,30^. In panels B and C, ***p<0.001, **p<0.01, *p<0.05, and n.s. p>0.05.

We first assessed how variability affects learning itself. To this end, we computed the “Learning Index” (LI) ^28^, which quantifies learning by comparing performance on the last (fourth) training day to baseline performance. The average improvement in LI was 0.13 (Fig. 1B; low variability mean LI=0.13, SE=0.02, permutation test against 0, p<0.001, Hedges’ g=1.95; high variability mean LI=0.13, SE=0.03, permutation test, p<0.001, Hedges’ g=1.55), and there were no significant differences in LIs between the groups (mean difference in LI=0.008, SE=0.03, permutation test, p=0.815, Hedges’ g=0.07). This suggests that SF variability did not negatively affect the amount of PL in the orientation discrimination task.

We obtained similar results when we quantified perceptual thresholds instead of LIs (Fig. 1C): initial thresholds on day 1 did not differ between the high and the low variability groups (mean difference=1.41°, SE=0.855°, permutation test, *p=*0.105, Hedges’ *g=*0.52). This suggests that variability did not negatively affect task performance at the outset of training. On average, learning resulted in a decrease of orientation discrimination thresholds by 2° in both training groups (low variability [last-first]=−2.576°, SE=0.623, permutation test against 0, p<0.001, Hedges’ g=−1.08; high variability [last- first]=−1.43°, SE=0.584, permutation test, p=0.033, Hedges’ g=−0.67). Threshold improvements did not differ significantly between the groups (mean difference between groups [last-first]=1.15°, SE=0.854, permutation test, p=0.197, Hedges’ g=0.43).

To test generalization, on the fifth day, we changed the stimulus location to a new isoeccentric (12.4°) location (8 dva away, upper right quadrant, see Methods). As a result, we found a significant drop in discrimination ability (i.e., increase in threshold) for the low variability group (mean threshold increase [transfer-last]=0.62°, SE=0.350, permutation test, p=0.039, Hedges’ g=1.08). No such effect was evident in the high variability group (mean threshold increase [transfer1-last]=0.21°, SE=0.564, permutation test, p=0.361, Hedges’ g=0.09). Together, these behavioral results suggest that task-irrelevant variability during a high-precision orientation discrimination task indeed enables generalization in PL, while low variability leads to specificity, replicating our previous findings ^12^.

### Changes in early visual representations

Building on the behavioral findings, we next investigated the neural substrates underlying PL and how they account for the observed transfer effects. We first focus on changes in early visual representations at the retinotopically defined training locations in areas V1, V2, and hV4, which were independently localized using standard retinotopic mapping procedures. If the brain flexibly identifies a neural “sweet spot” that depends on the training regime, we hypothesized that orientation training with low SF variability would drive learning within early visual areas (e.g., V1, V2), which contain finely tuned neurons with small receptive fields, whereas training with high variability would engage more SF-invariant populations in higher-order visual areas (e.g., hV4).

To test this hypothesis, we used MVPA and trained a classifier to discriminate clockwise from counterclockwise orientations using the spatial pattern of fMRI blood oxygenation level-dependent (BOLD) activity at the training location (Fig. 2A, see Methods). Learning, quantified as the change in orientation information between training days 1 and 4, was similar across all three regions of interest (ROIs) within early visual cortex (EVC; Fig. 2B), with strong positive correlations between learning effects in V1, V2, and hV4 (all r ≥ 0.93, p<0.001, FDR-corrected). In particular, we used a linear mixed-effects model (LME) including the factors Session, Group, and ROI (see Methods). Neither the full three-way interaction (ΔAIC=12, χ²(7)=2.00, p=0.96) nor the Session × Group interaction (ΔAIC=0.76, χ²(1)=1.23, p=0.27) were significant (all p>0.267) nor improved the model fit, hence, results are reported from the main-effects-only model. This model revealed a significant learning effect (Session: β=0.046, SE=0.012, t(235)=3.81, p<0.001), but no effects of Group (β=−0.008, SE= 0.019, p=0.69) or ROI (all ps>0.85). The average increase in decoding accuracy between sessions was 4.64% (SE=1.22%). Hence, while orientation representations improved with training, this improvement did not differ between training regimes, suggesting that differences in generalization cannot be explained by representational changes in EVC.

**Figure 2:**
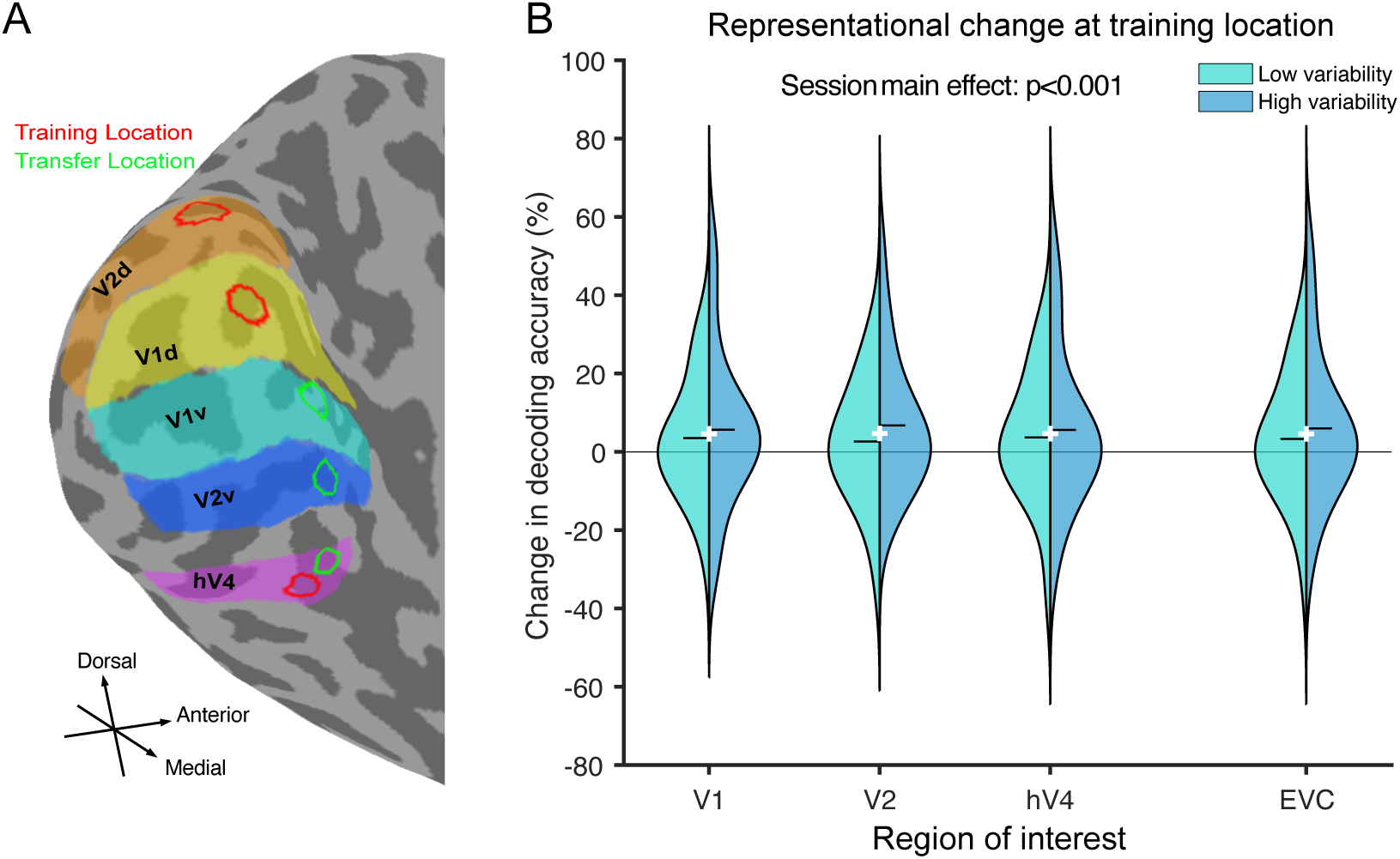
MVPA decoding in early visual cortex. **A.** Within early visual cortex (EVC), we defined three ROIs corresponding to the training location (red circles) and three corresponding to the transfer location (green circles) using retinotopic mapping. As shown in this example left hemisphere (subject 09, the selected areas included V1, V2, and hV4, subdivided into V1 dorsal (V1d, yellow), V1 ventral (V1v, pale blue), V2 dorsal (V2d, orange), V2 ventral (V2v, dark blue) and hV4 (violet). **B.** Violin plots show the change in decoding accuracy of a MVPA classifier trained to distinguish clockwise from counterclockwise orientations based on single-trial beta estimates obtained with the LSS procedure. Decoding accuracy improved across sessions (Session main effect=0.046, SE=0.012, t(235)=3.81, p<0.001), consistent with PL, but no significant differences were observed between groups (β=–0.008, SE=0.019, t(235)=–0.40, p=0.69) or ROIs (all ps>0.85). The average increase in decoding accuracy across groups and ROIs from the first to last session was 4.64% (SE=1.22%). Each violin plot depicts the kernel-density distribution of the data per group. The width corresponds to the relative frequency of observations, the central bar marks the respective group’s median, and the white cross indicates the mean over groups.

### Changes in behaviorally relevant functional connectivity

Alternatively or in addition to changes in representations themselves, PL may also affect the connectivity between brain regions, which has been taken as a proxy for readout of visual information, e.g., by decision making circuits ^26^. To assess whether PL indeed affected inter-regional connectivity, and whether such connectivity changes were modulated by the training regime, we quantified behaviorally relevant functional connectivity by means of psycho-physiological interaction (PPI) analysis. Specifically, we investigated whether functional connectivity that was differentially modulated on correct versus incorrect trials changed with PL in the two training groups. We focused on connectivity between the retinotopic stimulus locations in early visual areas and intraparietal sulcus (IPS) subregions involved in perceptual decision-making (Fig. 3A). To this end, we predicted the time courses of each target ROI (IPS0, IPS1–3, IPS4–5), respectively, using LME models with fixed factors for each seed ROI time course (V1d, V2d, or hV4), behavioral accuracy, session, group, and their interactions (whereby the interaction of the seed time course with the behavioral predictor correct/incorrect, i.e., the PPI predictor, reflects behaviorally relevant connectivity, see Methods). This approach allowed us to quantify connectivity changes over learning (PPI × Session) and directly test the effects of training variability (PPI × Session × Group). Indeed, we found learning effects that were common among groups (Fig. 3B) as well as learning effects that differentiated high from low variability training (Fig. 3C): the earliest visual cortex seed region to show a learning effect on connectivity was V2d, which exhibited a significant increase in connectivity from the first to the last training session with IPS0 (PPI × Session [last–first]=0.021, SE=0.0085, t(134380)=2.49, p=0.013, FDR-corrected p<0.05). These learning-related changes did not differ between groups, as all PPI × Session × Group interactions were non-significant (all ps>0.25), and no other V2d–IPS connections (IPS1–3, IPS4–5) showed significant effects (all ps>0.38). Hence, both groups increased functional connectivity between V2 and a posterior IPS region during PL. In contrast, V1d showed no significant learning effect on connectivity (PPI × Session, all ps>0.25) and no effects related to training variability (all ps>0.12) for either group.

**Figure 3.**
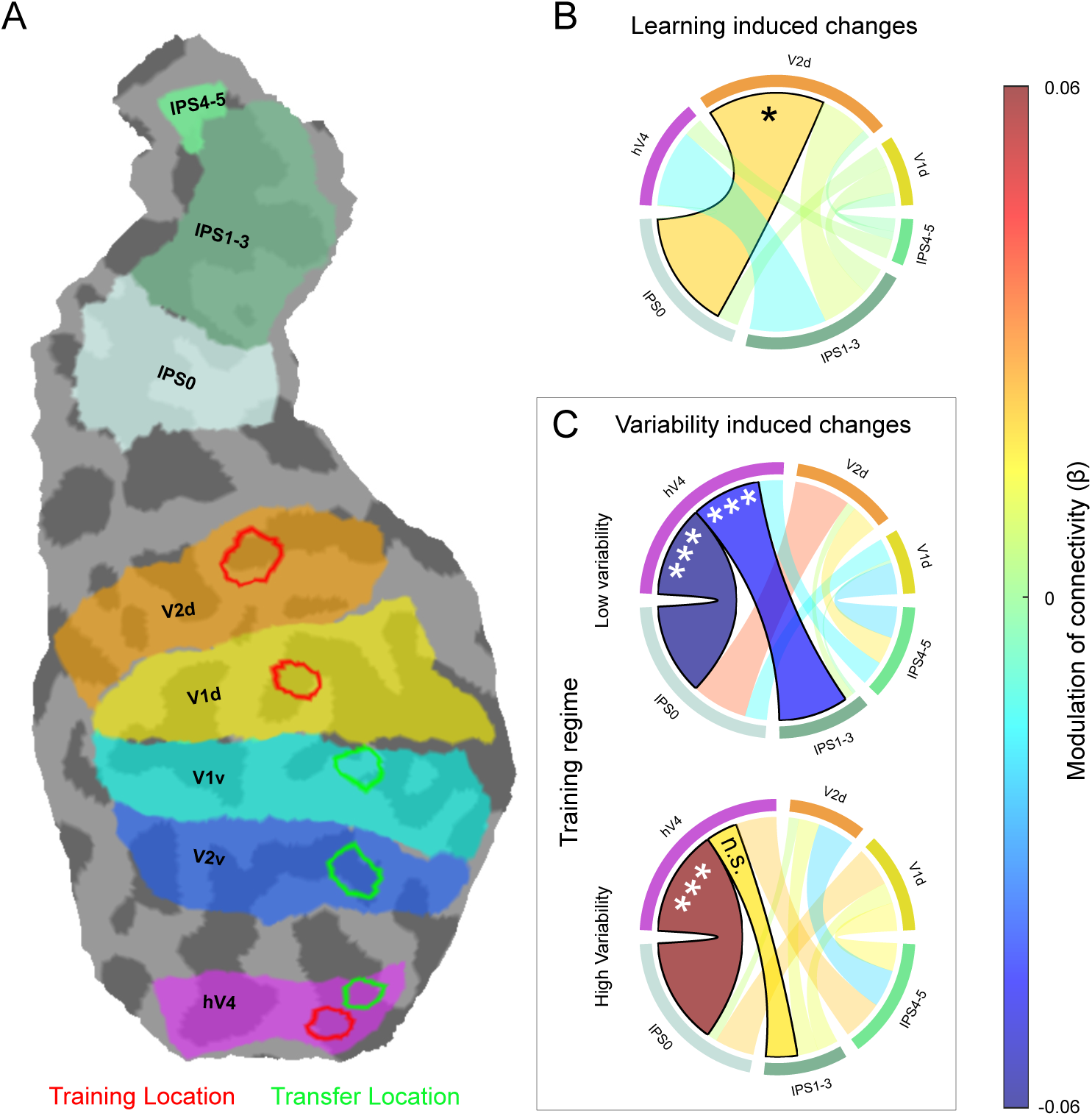
Connectivity results: session and variability effects. **A**. Flat map of an example hemisphere displaying (subject 09 three ROIs at the training location (red) and three at the transfer location (green), defined using individual retinotopic maps. Seed regions included V1, V2, and hV4, subdivided into V1 dorsal (V1d, yellow), V1 ventral (V1v, pale blue), V2 dorsal (V2d, orange), V2 ventral (V2v, dark blue), and hV4 (violet). Target regions were defined within the intraparietal sulcus (IPS) as IPS0 (light green), IPS1–3 (sea green), and IPS4–5 (spring green). **B.** Training induced a significant increase in V2d–IPS0 connectivity across sessions irrespective of training regime (PPI × Session [last–first]=0.021, SE=0.0085, t(134,380)=2.49, p=0.013, FDR-corrected p<0.05). No other V2d–IPS connections (IPS1–3, IPS4–5) showed significant changes (all ps>0.38). **C.** Effects of variability on connectivity were found between hV4 and IPS0 (PPI × Session × Group [low-high, last-first]= –0.120, SE=0.022, t(134,380)=–5.36, p=8.4 × 10⁻⁸, FDR-corrected p<0.001), and between hV4 and IPS1–3 (PPI × Session × Group [low-high, last-first]=–0.064, SE=0.025, t(134,380)=–2.60, p=0.009, FDR-corrected p<0.05). Low SF variability led to a decrease in connectivity between hV4 and IPS0 (low variability; PPI × Session [last-first]=–0.060, SE=0.016, t(134,380)=−3.77, p<0.001, FDR-corrected p<0.001) and IPS1–3 (low variability [last-first]=–0.046, SE=0.017, t(134,380)=−2.63, p=0.008, FDR-corrected p<0.05), whereas high variability induced significant increases in connectivity between hV4 and IPS0 (high variability [last-first]=+0.060, SE=0.016, t(134,380)=3.81, p<0.001, FDR-corrected p<0.001). In all panels, significant changes in connectivity are highlighted with thicker connection borders. In all panels, p-values are FDR-corrected: ***p<0.001, **p<0.01, *p<0.05, n.s. p>0.05.

A different picture emerged for the hierarchically highest retinotopically organized visual area we considered, hV4 (Fig. 3C). Here, we found a robust effect of training variability: specifically, connectivity between hV4 and IPS0 differed significantly across training regimes (PPI × Session × Group [low-high, last-first] =–0.120, SE=0.022, *t*(134380)=–5.36, *p*=8.4 × 10⁻⁸, FDR-corrected *p*<0.001), as did connectivity between hV4 and IPS1–3 (PPI × Session × Group [low-high, last-first] =–0.064, SE=0.025, *t*(134380)=–2.60, *p*=0.009, FDR-corrected *p*<0.05). Notably, the group-specific connectivity changes revealed opposite trends for the two training conditions: in the low-variability group, connectivity decreased significantly between hV4 and IPS0 (low variability; PPI × Session [last-first]=–0.060, SE=0.016, *t*(134380)=−3.77, *p*<0.001, FDR-corrected *p*<0.001) and between hV4 and IPS1–3 (low variability [last-first]=–0.046, SE=0.017, *t*(134380)=−2.63, *p*=0.008, FDR-corrected *p*<0.05). In contrast, the high-variability group showed increased connectivity both between hV4 and IPS0 (high variability [last-first]=+0.060, SE=0.016, *t*(134380)=3.81, *p*<0.001, FDR-corrected *p*<0.001), but not between hV4 and IPS1–3 (high variability [last-first]=+0.018, SE=0.017, *t*(134380)=1.04, *p*=0.2992, FDR-corrected *p*>0.05). No significant variability-related training effects on connectivity were observed between hV4 and IPS4–5 for either group (all FDR-corrected ps>0.12).

Together, these results suggest that PL reshapes visual–parietal interactions, with V2d–IPS0 connectivity reflecting practice-related reweighting irrespective of the training regime, and task-irrelevant variability selectively modifying hV4–IPS coupling.

To test whether the variability-related increases in hV4–IPS connectivity also explicitly support generalization, we examined PPI effects in the transfer condition, where the stimulus was presented at a new location. Based on group-specific training effects between hV4d and IPS0 and IPS1–3, respectively, we assessed how these regions communicate during the transfer session. Using accuracy-based PPI and a group contrast (low vs. high variability), we observed a significant effect for hV4v–IPS0 connectivity (PPI × Group [low–high]=–0.029, SE=0.0128, t(67177)=–2.25, p=0.025, FDR-corrected p<0.05), reflecting stronger coupling in the high variability than in the low variability group. hV4v–IPS1–3 connectivity showed no significant group differences (PPI × Group [low–high]=0.0017, SE=0.0124, t(67177)=0.14, p=0.89, FDR-corrected p>0.05). These results further support a differential role of hV4-IPS coupling in enabling variability-induced generalization of orientation discrimination PL to new spatial locations.

## DISCUSSION

In this study, we investigated how task-irrelevant variability shapes PL and its generalization. Behaviorally, both low and high variability training produced comparable improvements in the trained orientation discrimination task (Fig. 1). Critically, high variability training enabled transfer to a novel stimulus location, while low variability training led to specificity, replicating prior evidence that task-irrelevant variability promotes generalization ^12,17^. At the neural level, PL improved orientation representations in early visual areas (V1, V2, hV4), but no differences emerged between training groups (Fig. 2B). This indicates that introducing task-irrelevant variability during training does not alter representations in early visual cortex and that the balance between specificity and generalization cannot be explained by local sensory plasticity alone. Instead, our findings point to network-level mechanisms as the substrate for generalization. Functional connectivity analyses revealed a common learning-related increase in connectivity between V2 and IPS0 across groups (Fig. 3B), consistent with a shift toward reweighting sensory representations at the decision stage ^9^. Crucially, high variability training relied on task-dependent connectivity between more SF-invariant area hV4 and decision-related regions in IPS, whereas low-variability training decreased connectivity along the same pathway. Together, these results demonstrate that the brain flexibly engages distinct learning configurations depending on the training environment, giving rise to either specific or generalizable learning (Fig. 3C).

Our findings align with classical theories of PL but extend them by showing that the neural locus of learning is not fixed; rather, it flexibly adapts to the structure of the training environment. Low variability training reproduced the classical signature of specificity ^18,20,31^, engaging early visual areas with small receptive fields and sharp tuning, thus confining improvements to the trained feature and location. High variability training enabled transfer to untrained locations and recruited higher-level, SF-invariant visual areas with larger receptive fields. Together, these results support a “sweet spot” model of PL, in which the training regime determines the neural substrate that supports behavioral improvement (Fig. 4).

**Figure 4:**
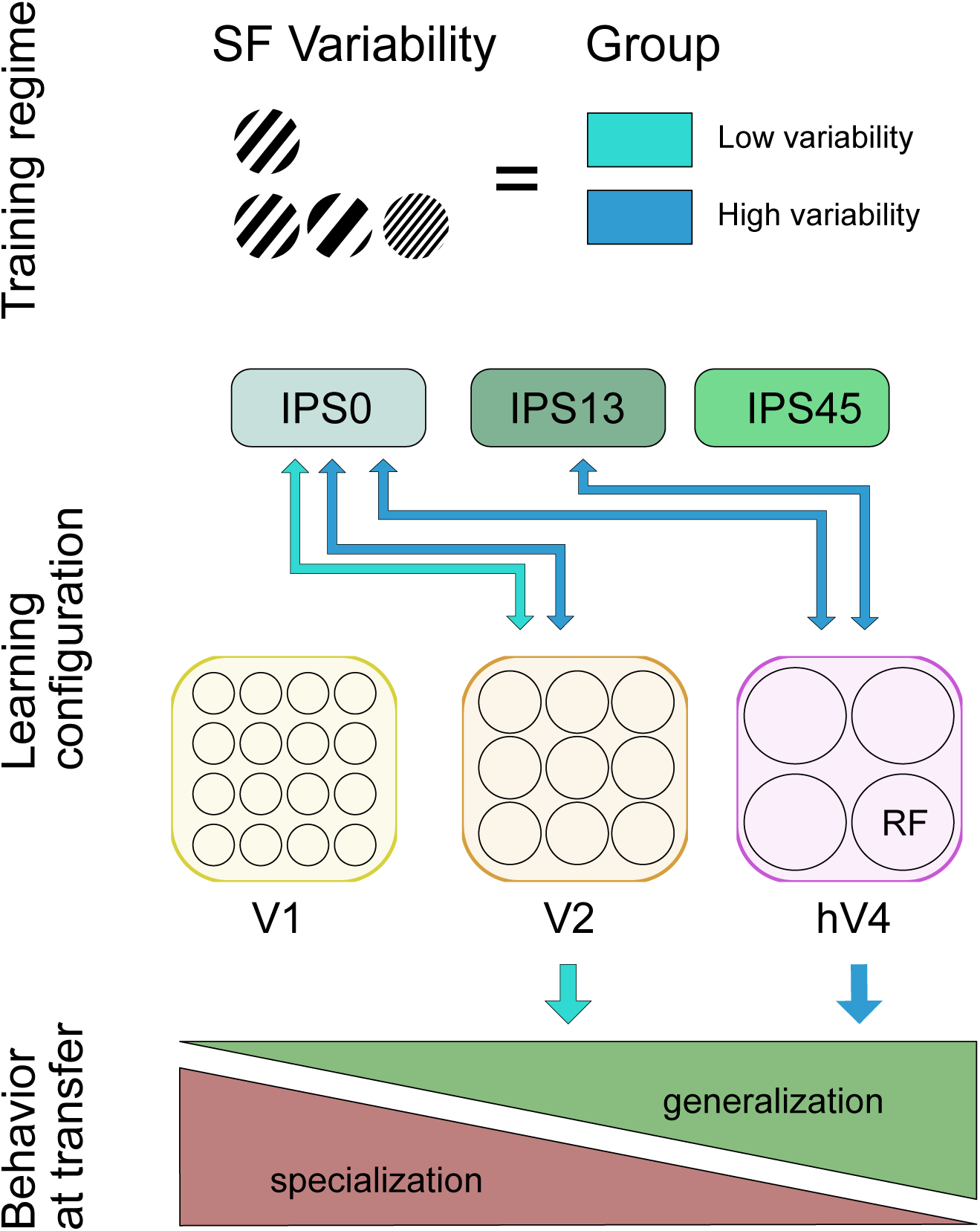
The “sweet spot” hypothesis of PL: The brain appears to adopt alternative strategies depending on the training environment, leading to distinct neural implementations of learning and, consequently, different patterns of transfer. Perceptual learning may thus emerge from distinct physiological “sweet spots” across the sensory hierarchy. Training with high task-irrelevant SF variability promotes recruitment of SF-invariant neurons with large receptive fields that facilitate generalization across stimuli. In contrast, low variability training engages more narrowly tuned populations optimized for precision but restricted in transfer. As a result, when learning is tested with a new stimulus dimension, such as an untrained location, only the group relying on invariant neural populations shows generalization.

Our “sweet spot” model could help explain the diversity of findings in the electrophysiological literature through a common underlying principle: electrophysiological studies of PL show learning-related plasticity at multiple hierarchical stages, from early visual cortex ^25^, to higher areas such as V4 ^32–34^, and even TEO (Adab, Popivanov, Vanduffel, & Vogels, 2014). The diversity of reported loci across studies may therefore reflect differences in training dimensionality or variability ^12,36^, rather than contradictory neural mechanisms. Similarly, our results may resolve the question why the causal contribution of brain areas to behavior changes depending on training history ^37^, i.e., hierarchically higher visual areas causally contribute to behavior after training with a more diverse stimulus set than after training with only one stimulus dimension. Future studies could test this hypothesis in humans using causal interventions, such as transcranial magnetic stimulation, to selectively modulate different stages of hierarchically processing and determine their contribution to generalization in PL.

Our data supports the view that PL depends not solely on local sensory plasticity but also on how higher-order regions read out information from visual cortex. Learning-related changes in task-relevant functional connectivity thus offer a parsimonious mechanism, consistent with adjustments to effective feedforward weights ^26^ (although our design does not allow us to rule out contributions of feedback signals). The “sweet spot” of PL emerged in connectivity changes between visual cortex and IPS regions: posterior IPS0 and mid IPS1-3 are tightly coupled to retinotopic visual areas ^38–40^; in combination with large receptive fields ^41^, they enable spatially broad pooling and naturally account for generalization beyond the trained quadrant we observed behaviorally. Anterior IPS4–5 shows minimal visual connectivity and stronger coupling with prefrontal and premotor areas ^38,40^, making it an unlikely site for early sensory readout optimization. Importantly, for the “sweet spot” hypothesis - which rests on invariance as the basis for generalization - it is immaterial whether plasticity occurs within representations or within the connectivity that governs their readout, as both yield the same functional outcome.

Our results have several broader implications. They offer a mechanistic account of how variability in training promotes generalization, highlighting that learning is most effective when cortical circuits operate in a regime that supports invariant readout. By linking behavioral improvements to changes in task-relevant connectivity, the findings point to an interaction between local sensory plasticity and downstream readout processes, consistent with whole-brain perspectives on PL ^42,43^. More broadly, they demonstrate the flexibility of cortical networks in adopting alternative computational strategies depending on training structure. The resulting “sweet spot” principle may help inform rehabilitation, perceptual expertise training, and educational approaches where generalization beyond the trained conditions is essential ^44^.

## RESOURCE AVAILABILITY

### Lead contact

Further information and requests for resources and reagents should be directed to and will be fulfilled by the lead contact, Caspar M. Schwiedrzik.

### Materials availability

This study did not generate new unique reagents.

### Data and code availability

- All human behavioral data will be made available for download on Figshare upon acceptance.
- Any to re-analyze the data reported in this paper is available from the lead contact upon request.

## ACKNOWLEDGMENTS

We would like to thank Peter Dechent, Carsten Schmidt-Samoa, Britta Perl, Ilona Pfahlert, Daniela Bernd, and Kristin Kötz for help with data acquisition. This project has received funding from the European Research Council (ERC) under the European Union’s Horizon 2020 research and innovation program (Grant agreement No. 802482, to CMS). C.M.S. is supported by the German Research Foundation’s Emmy Noether Program (SCHW1683/2-1). The funders had no role in study design, data collection and interpretation, decision to publish, or preparation of the manuscript.

## AUTHOR CONTRIBUTIONS

Methodology, investigation, formal analysis, visualization, data curation, writing – original draft preparation, G.L.M.; conceptualization, methodology, writing – original draft preparation, supervision, project administration, funding acquisition, C.M.S.

## Declaration of generative AI and AI-assisted technologies

In preparing this report, generative artificial intelligence (AI) tools were used solely to streamline the text. Specifically, ChatGPT was employed to improve clarity and coherence. No citations, references, or factual content were generated by AI.

## STAR METHODS

### KEY RESOURCES TABLE

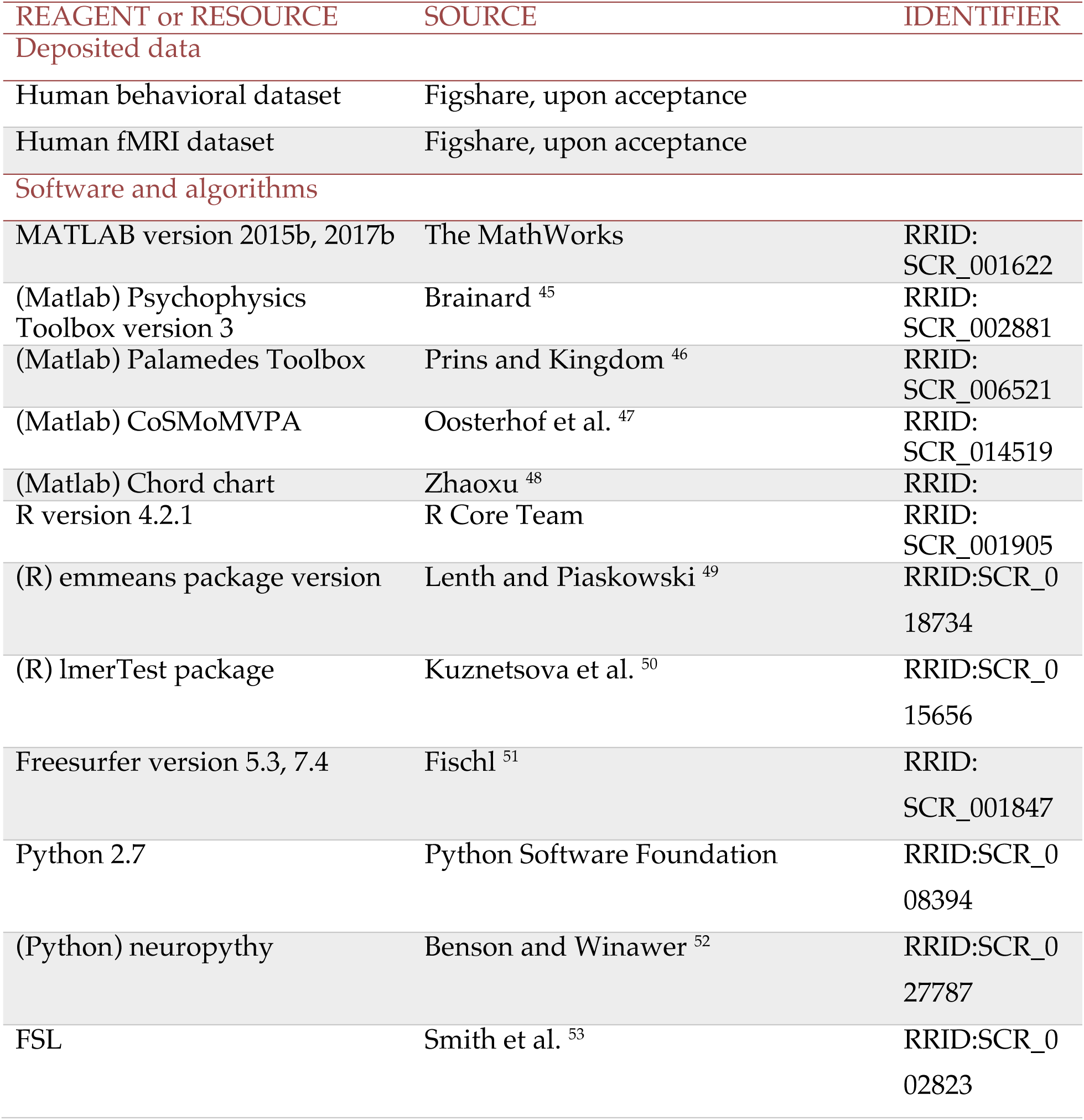

### EXPERIMENTAL MODEL AND SUBJECT DETAILS

#### Human Participants

A total of 52 healthy human volunteers (31 female, 4 left-handed, mean age 27 yrs, SD 6.5 yrs) participated in this study: All subjects had normal or corrected-to-normal vision, no neurological or psychiatric disease, and gave written informed consent before participation in accordance with the Declaration of Helsinki. The sample size was selected based on previous studies (Jehee, Ling, Swisher, Van Bergen, & Tong, 2012). Subjects were randomly assigned to one of two training groups (crossing the factor variability (high/low), see below). All procedures were approved by the Ethics Committee of the University Medical Center Göttingen (protocol number 29/8/17).

### METHOD DETAILS

#### General setup

All subjects were trained on a two-alternative forced choice (2AFC) orientation discrimination task with oriented gratings. Training lasted 4 days with one training session per day, followed by a transfer session. Total training and transfer time was 5 days.

The first training, last training, and transfer days were performed within the MRI setup. In this setup, stimuli were projected on an MR-compatible projection screen (43cm x 24 cm, visible range 27.2° horizontally and 15.4° vertically) via a ProPixx projector (Viewpixx, refresh rate 120, resolution 1920×1080 pixel, viewing distance to participants’ eyes 89 cm) at the end of the bore of a 3T MRI scanner (Prisma, Siemens Healthineers). Stimulus delivery and response collection were controlled using Psychtoolbox ^45^ running in Matlab (The Mathworks, Inc.). Auditory feedback was delivered via MR-compatible earphones (Sensimetric S15, Sensimetrics Corporation) combined with an ear tip (Comply Foam Canal Tips) to maintain an acoustic seal and reduce MRI noise. During all experiments, we continuously acquired pupil and gaze measurements using an MR-compatible, mirror-mounted eye tracker (ViewPoint Eytracker, Arrington Research). Data were sampled at 60 Hz from both eyes. Subjects responded using two MR-compatible response boxes, one held in each hand. Left and right choices were indicated by button presses on the left- and right-hand response boxes, respectively. The built-in Siemens pulseoximeter was used during each scan session to monitor blood oxygenation (data not reported). Subjects were paid €15 per hour. To ensure constant motivation over the training sessions, subjects received a bonus of €2 if they improved by 10% from the previous training session.

Two training sessions (day 2 and day 3) were conducted outside the MRI scanner in our psychophysics setup. Stimuli were presented on an LCD monitor (ViewPixx/EEG, refresh rate 120 Hz, resolution 1920 × 1080 pixel, viewing distance 65 cm) in a darkened, sound-attenuating booth (Desone Modular Acoustics). Stimulus delivery and response collection were controlled using Psychtoolbox running in Matlab. Auditory feedback was delivered via headphones (Sennheiser HDA 280). During all experiments, we continuously acquired pupil and gaze measurements using a high-speed, video-based eye tracker (SR Research Eyelink 1000+). Data were sampled at 1000 Hz from both eyes. To keep the response modality identical between MRI and psychophysics sessions ^54^, participants used two response boxes, one held in each hand, with left- and right-hand button presses indicating left and right choices, respectively. Subjects were paid €12 per hour. To ensure constant motivation over the training sessions, subjects received a bonus of €2 if they improved by 10% from the previous training session.

#### Stimuli and task

On each trial, subjects had to decide whether a monopolar, monochromatic Gabor grating (size 3.1 dva, luminance 116.5 cd/m^2^ and 43.4 cd/m^2^ for the MRI and Psychophysics setup, respectively) was tilted clockwise or counterclockwise with respect to a reference stimulus. The reference stimulus was a monopolar, monochromatic Gabor grating with identical size and constant spatial frequency (SF, 2.56 cpd), tilted 27° from the horizontal meridian. The task stimuli were presented in five linearly spaced difficulty levels clockwise and counterclockwise from the reference, respectively. Each condition was presented 28 times. In addition, we presented the reference orientation 28 times, amounting to a total of 308 trials per session evenly distributed among four runs. Difficulty levels ranged from 0.5 to 2.75 deg deviation from the reference stimulus. Task-irrelevant variability was controlled as in our previous study ^12^. In the low variability group, all stimuli were presented at a single SF (1.70 cpd). In the high variability group, we used three SFs (0.53, 1.70, and 2.76 cpd). We chose these SFs such that they lie outside the other spatial frequency channels (including the one used for the transfer, see below), or exactly at full width half maximum (FWHM), with the exception of 1.7 cpd which just falls into the 2.76 cpd band. For this, we assumed a SF channel bandwidth of 1.4 octaves ^55^ (but see ^56^). Note that although SF and orientation are not independently processed in the visual system, we rendered SF task irrelevant by instructing subjects to consider only orientation for the task. In all groups, stimuli were presented in pseudo-random order at 12.4° eccentricity against a grey background (105.9 cd/m^2^ and 39.5 cd/m^2^ for the MRI and psychophysics setup, respectively). Phase varied randomly between 0° and 360° from trial to trial.

On each run, we first presented the reference stimulus for 5000 ms. A given trial started with two grey saccade placeholders (luminance 63.1 cd/m^2^ and 23.5 cd/m^2^ for the MRI and psychophysics setup, respectively) placed at 12 dva horizontal distance from the center of the screen and a black fixation point constantly displayed on the screen. This was followed by a 1000 ms fixation period, followed by the stimulus for 250 ms. After a pseudorandomly distributed delay of 500-20000 ms, the choice phase started, which was indicated by a color change of the placeholders. The red target (luminance 63.1 cd/m^2^ and 23.5 cd/m^2^ for the MRI and psychophysics setup, respectively) was the instructed target to choose in order to report that the stimulus was rotated counterclockwise, while an isoluminant green target indicates it was rotated clockwise. To respond, subjects had to press a button on the same side as the colored target: e.g., to report clockwise, if the green target was on the right, the subject had to press a button on the right. The assignment of colors to the placeholder locations was pseudo-randomized. Subjects had to choose the target within 1000 ms. Subjects were instructed to respond as accurately as possible. Feedback on accuracy was provided by playing a low-pitch sound (incorrect) or a high-pitch sound (correct) for 500 ms. The next trial started after a pseudorandomly distributed inter-trial interval of 500-20000 ms. If subjects did not respond in time or if they broke eye fixation (fixation window size 4.5 dva), the low-pitch sound was played, and the trial was repeated later during the block. To keep subjects motivated, the high pitch sound increased in loudness after the first 2 and 3 sequential correct trials. Loudness was reset to the original level at the first incorrect trial. All runs had a constant duration of 630000 ms, on average each trial could last a maximum of 7400 ms allowing subjects to perform all 77 trials plus 8 repetitions.

At the end of each run, feedback about the percent correct performance was provided to the subjects and they were allowed to take a short break. As a reminder, the reference was shown for 5000 ms at the center of the screen before each block (in total 4 times per session). Before starting the first experimental session, subjects were familiarized with the task with several warm-up trials.

#### Transfer condition

For the transfer test, we changed the location of the stimuli to a new, iso-eccentric position in the upper quadrant 8 dva away from the original training location (lower quadrant), similar to previous studies ^12,17,57^. All other parameters remained identical to the training conditions. Stimulus SF was kept identical as during training. Subjects again performed 308 trials within a single session.

#### Retinotopic mapping

Retinotopic mapping was carried out using a rotating checkered wedge or an expanding checkered ring, respectively ^58,59^. For eccentricity mapping, 20 loops of expanding checkered concentric rings were displayed. The rings expanded from the fixation point to the maximal eccentricity of 16 dva. Polar angle was mapped with a rotating checkered wedge, which covered ¼ of the screen (20 loops). Checkerboards inverted polarity at a rate of 8Hz. Subjects were instructed to fixate on the fixation dot until the end of the measurement.

#### MRI data acquisition

MRI data were collected with a 64-channel head coil on a 3T scanner (Prisma, Siemens Healthineers). Anatomical images were acquired using a Magnetization-Prepared 2 Rapid Acquisition Gradient Echoes (MP2RAGE) T1-weighted sequence (FOV: 256 x 256mm; voxel size: 1 x 1 x 1mm; TR: 5000ms; TE: 2.9ms; number of slices: 176) for the first and last session, while a shorter Magnetization-Prepared Rapid Acquisition Gradient Echo (MPRAGE) T1-weighted sequence (FOV: 256 x 256mm; voxel size: 1 x 1 x1mm; TR: 2250ms; TE: 3.3ms; number of slices: 176) was used for the second MRI session. Functional images for the main experiment were acquired using an echo-planar imaging (EPI) sequence (FOV: 210x 210mm; voxel size: 2 x 2 x 2mm; slice thickness: 2mm; multiband 3x, TR: 1500ms; TE: 30ms; 420 Volumes; flip angle: 70°; number of slices: 69). During MRI sessions 1 and 2, a 10 minutes retinotopic mapping was acquired (FOV: 256 x 256mm; voxel size: 2 x 2 x2mm; multiband 3x, TR: 1500ms; TE: 30ms; 210 Volumes; flip angle: 70°; number of slices: 69). In addition to the functional images, we recorded a short field correction scan before each run (with opposite phase encoding direction, for topup, see below).

#### Behavioral data analysis

Human behavioral data were analyzed using linear mixed effects (LME) models and (exact) permutation t-tests (two-sided unless otherwise noted). For t-tests, we computed Hedges’ g as the effect size. From the original sample of 52 subjects, 2 subjects did not complete the experiments and were thus excluded from data analysis. 5 subjects were excluded from further data analysis because they finished less than 50% of trials during the first session. 5 subjects were excluded during data acquisition because of a lack of significant learning (Learning Index = 0, see below). The final N in the main experiment was thus 40 (23 female, 3 left-handed, mean age 27 yrs, SD 6.0 yrs). 20 subjects (10 females, 1 left-handed, mean age 26.6 yrs, SD 2.2 yrs) were trained with 1 spatial frequency (low variability), and 20 subjects (13 female, 2 left-handed, mean age 27.6 yrs, SD 7.9 yrs) were trained with 3 different spatial frequencies (high variability group). Accuracy was defined as the average percentage correct per session. We excluded all trials with outliers in the reaction times per subject using the estimator Sn ^60^ at a threshold of 8.5, as in our previous study ^12^. This led to an exclusion of an of average 0.2% of trials per session. To quantify learning, we computed the Learning Index (LI), as in ^28^:

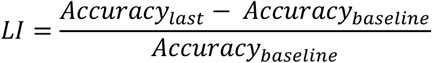

where baseline accuracy was obtained by averaging the performance of the first session. Because LI could not be lower than 0, we used one-sided (exact) permutation t-tests for comparisons against 0. To quantify transfer, we fitted psychometric functions (Weibull) using the Palamedes Toolbox ^46,61^ to derive orientation discrimination thresholds per subject and session.

#### MRI data analysis

##### Anatomical data analyses

Grey and white matter segmentation was obtained from the MP2RAGE and MPRAGE images using FreeSurfer ^51^ (http://surfer.nmr.mgh.harvard.edu/) via the recon-all pipeline as reported in ^62^. Before running the recon-all pipeline, the MP2RAGE images were denoised ^63^. Furthermore, to reduce the confounding effect of inter-individual morphological variability the reconstructed anatomical data were processed with the longitudinal-recon pipeline ^64^. This involved creating an unbiased within-subject template space and image from the multiple T1 measurements per subject ^65^, using robust, inverse consistent registration ^66^. This template informed subsequent processing steps like skull stripping, Talairach transformation, atlas registration, and generation of spherical surface maps and parcellations, leading to increased reliability and statistical power ^64^. Consequently, functional and anatomical images were aligned using boundary-based registration ^67^. Surface reconstruction was done following ^68,69^.

##### Functional data analyses

The functional data was preprocessed using FreeSurfer 5.3 and 7.4. We first performed motion correction ^70^ which performs rigid-body transformations on the images. In addition, we estimated and corrected the susceptibility-induced off-resonance field ^71^ using topup in FSL ^53^. To this end, we utilized images recorded with reversed phase-encode blips Functional data were smoothed with a 4 mm^3^ kernel and then projected onto the fsaverage surface space.

##### Regions of interest

Stimulus-driven ROI: To identify the retinotopic location of the stimulus for both training and transfer conditions within early visual areas, we used data from the retinotopic mapping scan. First, polar angle and eccentricity maps were computed using the FreeSurfer standard procedures in FS-FAST. These maps were then used to define the boundaries of V1, V2, V3, and hV4 following ^72^ and ^73^. Once these areal boundaries were obtained, stimulus-specific subregions within the ROIs were localized using a generalized linear model (GLM). Specifically, we applied a contrast comparing stimulus presentation in the lower quadrant (training location) versus the upper quadrant (transfer condition) from the polar angle mapping. In parallel, another contrast was applied to extract the portion of ROIs corresponding to the stimulus eccentricity of 12.4° from the eccentricity maps. The resulting significant surface vertices were combined through a conjunction ^74^. After visual inspection, all six resulting surface ROIs were scaled to the same size (3.5 mm) to ensure comparability across ROIs. This size was chosen because it represented the smallest common size across the six ROIs. Using this procedure, a subcluster of vertices was automatically defined within an approximately circular area, using the *–dilate-vertex* option of the FreeSurfer function *mri_binarize*.

For connectivity analyses, we defined IPS regions using the Wang atlas ^75^ for each participant based on anatomical templates provided by Benson and Winawer ^52^. Based on previous studies ^26,27^, our target regions were located within the intraparietal sulcus (IPS). We grouped the six atlas-defined IPS subregions into three clusters: IPS0, IPS1–3, and IPS4–5. These clusters broadly reflect distinctions between more visual (IPS0), intermediate (IPS1–3), and higher-order (IPS4–5) parietal subregions, as defined in prior work ^76–79^.

##### Multivariate pattern analyses

To quantify orientation representations, we used MVPA and trained a classifier to distinguish between clockwise and counterclockwise stimuli on the basis of BOLD activity. Classification was performed with a naïve Bayes algorithm (as implemented in CoSMoMVPA ^47^), using 50% of the trials for training and 50% for testing. For both training and test sets, we ensured a balanced distribution of labels. To maximize data usage and variability across trials, we applied the Least-Squares–Separate (LSS) procedure ^80,81^ to the preprocessed functional data using the FreeSurfer’s *selxavg3-sess*. This procedure produced single-trial beta estimates for each subject, session, and run (aligned to stimulus onset). A LME model was used to analyze changes in decoding accuracy across sessions, ROIs, and groups. The model included fixed effects for Session, ROI, and Group, as well as their interactions, and random intercepts for subjects. Among several candidate models of different complexity, the one reported here was selected as the best-fitting based on information criteria, providing an the best fit to the data (AIC=–400.13, BIC=–375.77, log-likelihood=207.07).

##### Psychophysiological interactions

We computed task-specific functional connectivity using a psychophysiological interaction (PPI) analysis ^82^. For each ROI, voxel time series were averaged and demeaned. The design matrix included the physiological timeseries averaged over the seed region, psychological regressors (accuracy: correct/incorrect, aligned to stimulus onset), group and session factors, and their interactions (up to four-way). Nuisance predictors, such as feedback time, choice onset, non-played time, and motion parameters including 12D motion correction, were also included. We then fitted a LME model across runs, sessions, subjects, and groups using R (lmer, lmerTest package), applying the Satterthwaite method for degrees of freedom. The model tested the effects of the seed ROI, psychological variables, and their interaction (PPI term) along with group and session factors. It also accounted for interactions among these variables, as well as the influence of nuisance predictors. Random intercepts were included for runs nested within subjects to account for repeated measurements. To interpret these effects, we extracted estimated PPI effects using the emtrends package ^49^. First, we assessed the session-dependent change in connectivity (PPI × Session) to capture learning-related effects common across groups. Second, we examined the interaction PPI × Session × Group to test whether connectivity changes across sessions differed between groups. Finally, we computed post hoc contrasts using a difference-of-differences approach to quantify group-specific changes in connectivity between sessions. We used the False Discovery Rate ^83^ procedure to correct for multiple comparisons across all target ROIs within each seed.

To identify PPIs during the transfer session, we followed the same preprocessing and design matrix structure used during the learning phase. Voxel time series within each ROI were averaged and demeaned. We then fitted a LME model, including fixed effects for the physiological regressor (seed ROI), the psychological regressor (correct/incorrect), PPI term, group, and their interactions, as well as nuisance predictors (see above). Random intercepts were included for subjects, with runs nested within subjects. To interpret connectivity effects, we estimated PPI slopes for each group using the emtrends package, and subsequently applied pairwise contrasts to test for group differences in connectivity during the transfer session. Chord charts shown in Figure 3B and C were created using a customized code adapted from ^48^. Both the width and color of the chords were linearly mapped to reflect the magnitude of connectivity modulation.

